# Spermiogenesis in the acoel *Symsagittifera roscoffensis*: nucleus-plasma membrane contact sites and microtubules

**DOI:** 10.1101/828251

**Authors:** Matthew J. Hayes, Anne-C. Zakrzewski, Tim P. Levine, Maximilian J. Telford

## Abstract

*Symsagittifera roscoffensis* is a small marine worm found in the intertidal zone of sandy beaches around the European shores of the Atlantic. *S. roscoffensis* is a member of the Acoelomorpha, a group of flatworms formerly classified with the Platyhelminthes, but now recognised as Xenacoelomorpha, a separate phylum of disputed affinity. We have used electron microscopy to examine the process of spermiogenesis (the final stage of spermatogenesis) in *S. roscoffensis*, by which spermatids form highly elongated spermatozoa. Their nuclei are long and thread-like, running most of the cell’s length and during the process a pair of flagella are fully incorporated into the cell body. Two previously undescribed inter-organelle contact sites form at different stages of spermiogenesis. Strikingly, there is an extensive nucleus-plasma membrane contact site. Golgi-derived granules containing electron-dense filaments line up along the spermatid plasma membrane, undergo a conformational change, and donate material that forms a peri-nuclear layer that cements this contact site. We also show in earlier stage spermatids that the same granules are associated with microtubules, presumably for traffic along the elongating cell. We identify a second spermiogenesis-specific contact site where sheaths engulfing each internalising flagellum contact the nuclear envelope. Finally, detailed studies of the spermatozoon axonemes show that the central keel has varying numbers of microtubules along the length of the cell, and is likely to be a centriole derivative.

**Summary sentence:** During spermiogenesis in the acoel flatworm *Symsagittifera roscoffensis,* two previously unidentified contact sites contribute to the structure of the mature spermatozoon and the axonemal structures show direct continuity between doublet and dense core microtubules.

## Introduction

*Symsagittifera roscoffensis* (von Graff 1891) (formerly *Convoluta roscoffensis*) is a small (5 mm long) acoel flatworm found inter-tidally on beaches in Europe. Adult individuals do not feed but rely on metabolites produced photosynthetically by a symbiotic alga, usually *Platymonas convolutae, Tetraselmis convolutae* or *Tetraselmis tetrathele* [1, 2]. The resulting green colour is the source of their common name of ‘mint-sauce worm’. The acoel flatworms are members of the Acoelomorpha, a group once thought to be part of the Platyhelminthes (true flatworms) but shown using molecular data to be an independent group whose true affinities remain controversial [3–9]. It is widely accepted that the Acoelomorpha (which contains acoels such as *Symsagittifera* and a second clade - the Nemertodermatida) are most closely related to the Xenoturbellida (another group of marine worms once associated with the Platyhelminthes) in a phylum called the Xenacoelomorpha. The position of the Xenacoelomorpha relative to other animal phyla is also controversial. Some molecular phylogenies place them as the sister-group to all other bilaterian phyla. This early emergence makes sense of the simple body plan of these worms. Other molecular studies conclude, however, that Xenacoelomorpha are related to the Ambulacraria (echinoderms and hemichordates) and that their morphological simplicity is a result of evolution by loss of characters that were present in a more complex ancestor.

Study of spermatozoa in invertebrates and in the acoels in particular has proven useful for comparative phylogenetic analysis [10–28]. Like many acoels, *S. roscoffensis* produce long, thread-like (filiform), spermatozoa which develop according to a process of ‘flagellar incorporation’ [29–31] (see schematics in Fig 1 and Fig 2). These sperm swim via undulations of a long tail containing two internalised, extended flagella and have undergone a ‘polarity inversion’, propelling the cell tail-first as compared to the head-first charge of archetypal ‘primitive’ forms. The mid-section of the typical acoel sperm contains many mitochondria and large granules and is supported by an axial axoneme bundle of dense-cored, atypical microtubules forming a so-called ‘keel’. These characteristics are believed to reflect the peculiarities of acoel fertilisation; the sperm often being introduced hypodermically into the body of the mate, penetrating between viscous cellular tissues as opposed to swimming freely in sea water as is the case with external fertilization. Although flagellar incorporation has been observed in numerous studies, a detailed mechanism of this process has not been proposed. As acoel sperm swim ‘tail first’ it has also been assumed by most that they do not have an acrosome, the large, granular vesicle that usually covers the crown of the nucleus in more ‘primitive’ aquasperm and which facilitates conjugation with the egg. These adaptions have called into question interpretation of them as representing the form of the ancestral bilateran.

**Figure 1.**
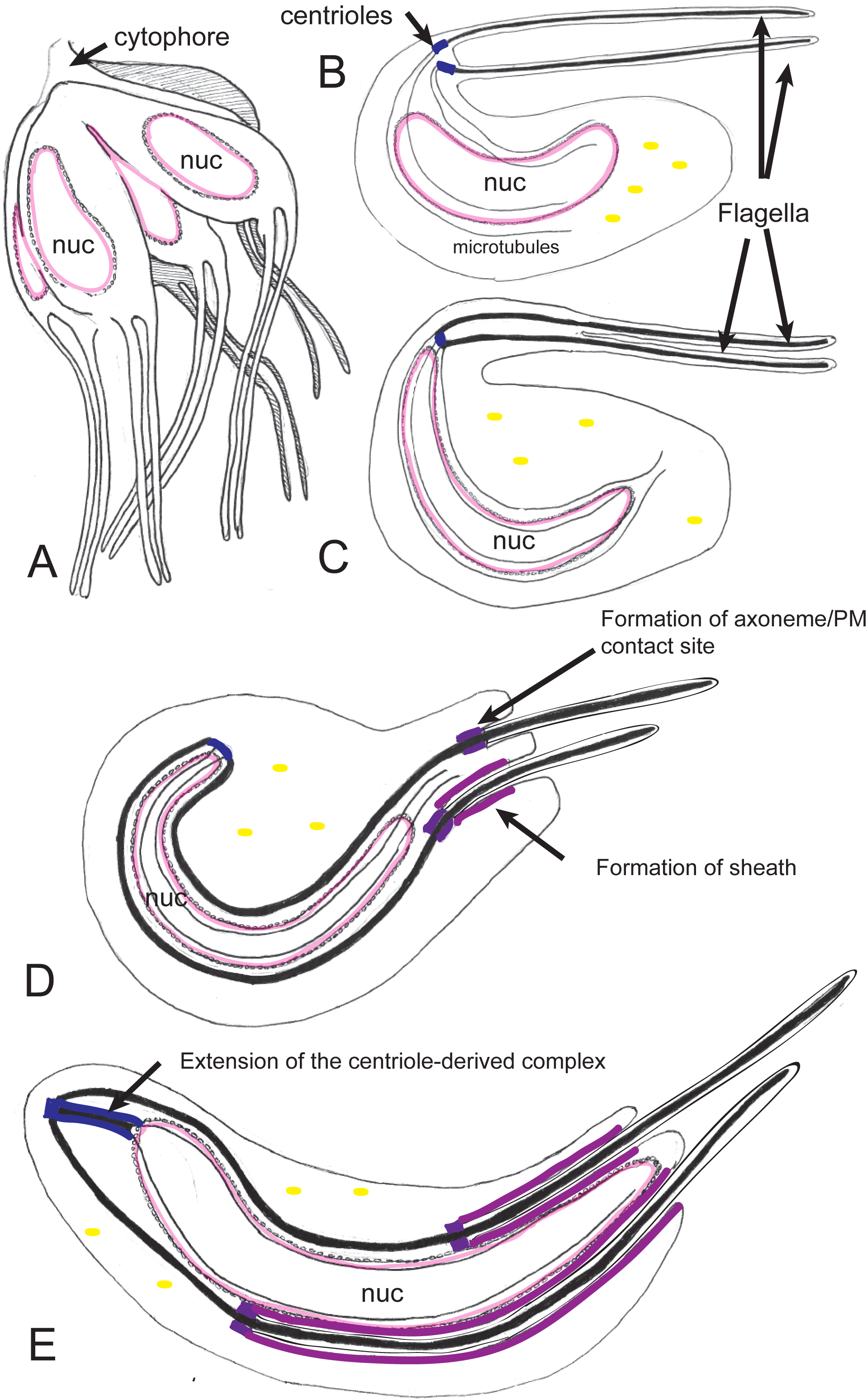
Schematic: Early stages of spermiogenesis. A: The products of meiosis remain closely associated by a cytophore c. B-C: (speculative) The basal bodies of the developing spermatocyte may have been separate or conjoined. They capture the nucleus; presumably via astral microtubules which wrap around the nuclear envelope (these giving rise to the manchette). D-E: The conjoined basal bodies and the internal elements of the flagella align with the elongating nucleus. The flagella begin to be assimilated into the cytoplasm of the spermatocyte, either by advancement of the cytoplasm over them, or by relative movements of the nucleus-associated axonemes and the cytoplasm. A multilaminate electron-dense complex and associated membrane contact site forms as the sheath develops, limiting a cytoplasmic canal, engulfing the flagella and linking them to the nucleus. E: The nucleus continues to elongate, as does the spermatocyte. The astral/manchette basal bodies remodel into the axial, atypical axoneme ’keel’, elongating and pushing the basal bodies away from the nucleus.

**Figure 2.**
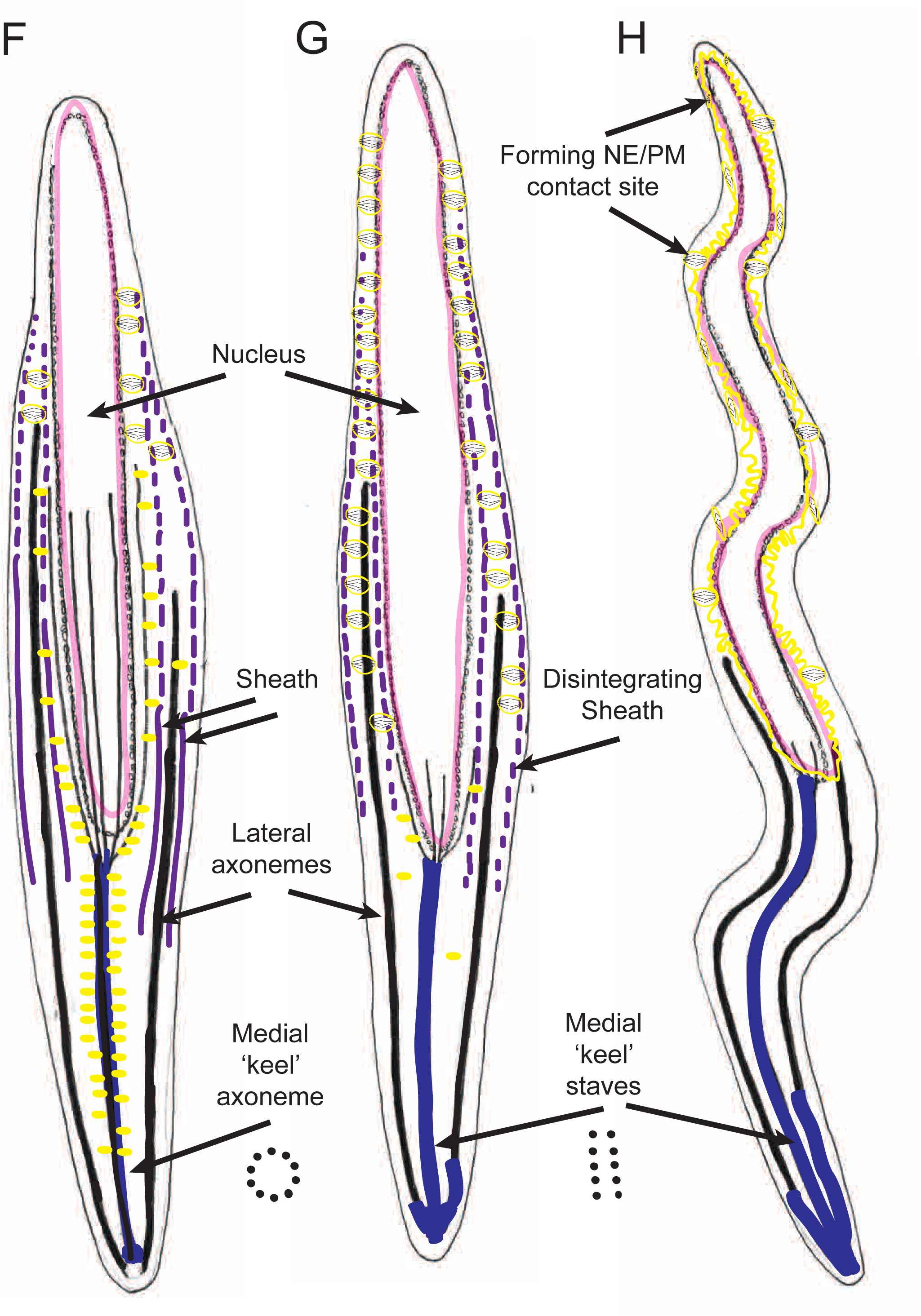
Schematic: Later stages of spermiogenesis. F: dense granules associate with the medial keel axoneme (a circular arrangement of atypical microtubules) and with the manchette (nuclear associated microtubules). These two elements are probably contiguous with one another. G: The medial keel becomes two staves; the dense granules lose their association with it and line up with the degenerating sheath and associated plasma membrane. The dense granules re-align their filamentous contents. H: The developing spermatozoan continues to elongate, the nuclear envelope becomes more elaborate as it merges with the dense granule membranes. Ultimately the filaments of the condensed granules will form a nuclear envelope/plasma membrane contact site.

In this paper we use transmission electron microscopy to examine spermiogenesis in *S. roscoffensis*. We identify previously overlooked aspects of the process, including the formation of unusual contact sites between membranes in the developing spermatozoon. Contact sites are specialised regions of organelle membranes that form close (usually less than 30nm, though there are exceptions) associations with one another. These sites seem to constitute dynamic mechanical tethers, holding the two organelles in position with respect to one another whilst organelle remodelling (structural and/or biochemical) occurs. They may be the site of transfer of ions, complex metabolites or lipids, and act as scaffolds for the recruitment of molecular machines required to act at the site of membrane-membrane association [38–39]. These contacts have been identified between nearly all organelles in most combinations. The ones we identify in this study occur during the process of flagella incorporation and also later during maturation of the complex nuclear envelope and give clues to mechanistic elements of spermiogenesis. This is the first description of an elongated nuclear envelope-plasma membrane contact site.

We have also examined in detail the axial axoneme, composed of electron-dense atypical microtubules, which appears to facilitate traffic of dense core granules that donate material for nuclear envelope-plasma membrane contact site formation. We have shown that the axial axoneme bundles distally with the internalised flagellar (lateral) axonemes. The ‘conventional’ doublets of the flagellar axonemes are contiguous with a ‘centriolar derivative’ composed of atypical microtubules identical to those of the axial axoneme. These observations, in broad agreement with other studies [16], suggest that the entire elongated axial axoneme is an extended centriole derived structure.

## Materials and Methods

### Specimens

Adult *Symsagittifera roscoffensis* were collected during low tide on the beach at Carantec (Brittany, France) in 2014. The animals were kept in plastic boxes with artificial seawater in incubators at 15°C with a 12h light/12h dark light cycle.

### Transmission electron microscopy

Adult specimens were fixed for two hours with cold (4°C) Karnovsky’s fixative (2% paraformaldehyde, 2.5% glutaraldehyde in 0.08M cacodylate buffer). They were washed 3 times in phosphate buffer and osmicated with 1% osmium tetroxide in ddH2O for 1 hour. Samples were then washed 3 x 10 minutes in ddH2O and dehydrated with a series of ethanol dilutions: 30%, 50%, 70%, 90%, 3 x 100% and 2 x propylene oxide (at least 20 minutes in each). They were infiltrated with 50:50 propylene oxide:araldite resin overnight and with several changes of 100% resin the next day. Blocks were cured at 60°C overnight. Sectioning was performed using a Leica Ultracut UCT microtome. Sections were counter-stained with Reynold’s lead citrate and were viewed on a JEOL 1010 TEM (JEOLUSA MA, USA).

### Preparation to enhance contrast in membranes (after Walton 1979 (40))

Specimens were fixed in Karnovsky’s EM fixative (as above) for 30 mins. They were washed 3 times with phosphate buffer. Samples were then incubated for 2 hrs in 2% osmium tetroxide/1.5% ferricyanide and then washed 3 x 5 mins in water. The samples were then placed in 1% thiocarbohydrazide solution for 10 minutes then washed 3 x 5 mins in water. A second round of osmication was performed (2% osmium tetroxide (no ferricyanide) for 30mins) and the tissue washed again 3 x 5 mins in water. The samples were then placed in aqueous 2% uranyl acetate overnight at 4 degrees.

The following day samples were placed in freshly made Walton’s lead aspartate (pH5.5) at 60°C for 30 mins and then washed 3 x 5 mins in water. Samples were then dehydrated through ethanol (30%, 50%, 70%, 90%) to 100% x 3 then placed in acetone 2 x 20 mins. Samples were infiltrated with Durcapan resin 25% plus acetone, then 50%, 75% (2 hours each) and 100% overnight. The following day the resin was exchanged, and the blocks hardened in an oven for 48 hours.

## Results

### Gross re-arrangements during spermiogenesis

Our general observations of spermiogenesis in *Symsagittifera roscoffensis* are in line with those of others who have used electron microscopy to examine related species (*Convoluta psammophila*) [11], (*Convoluta boyeri* and *Convoluta philippinensis*) [32–33], (*Paratomella rubra*) [17], (*Convoluta saliensis*) [34], (*Pseudaphanostema smithrii*, *Praeconvoluta tigrina*, *Convoluta pulchra*, and *Stromatricha hochbergi*) [26]. We observed multiple testes in various stages of development, each containing mature sperm and developing spermatocytes clustered around a *cytophore*, a residual domain of cytoplasm formed during meiosis that presumably serves as a communal source of biosynthesis (Fig 1A, Fig 3A). Individual spermatocytes at various stages of development and mature sperm were found in the same TEM sections in various orientations.

**Figure 3.**
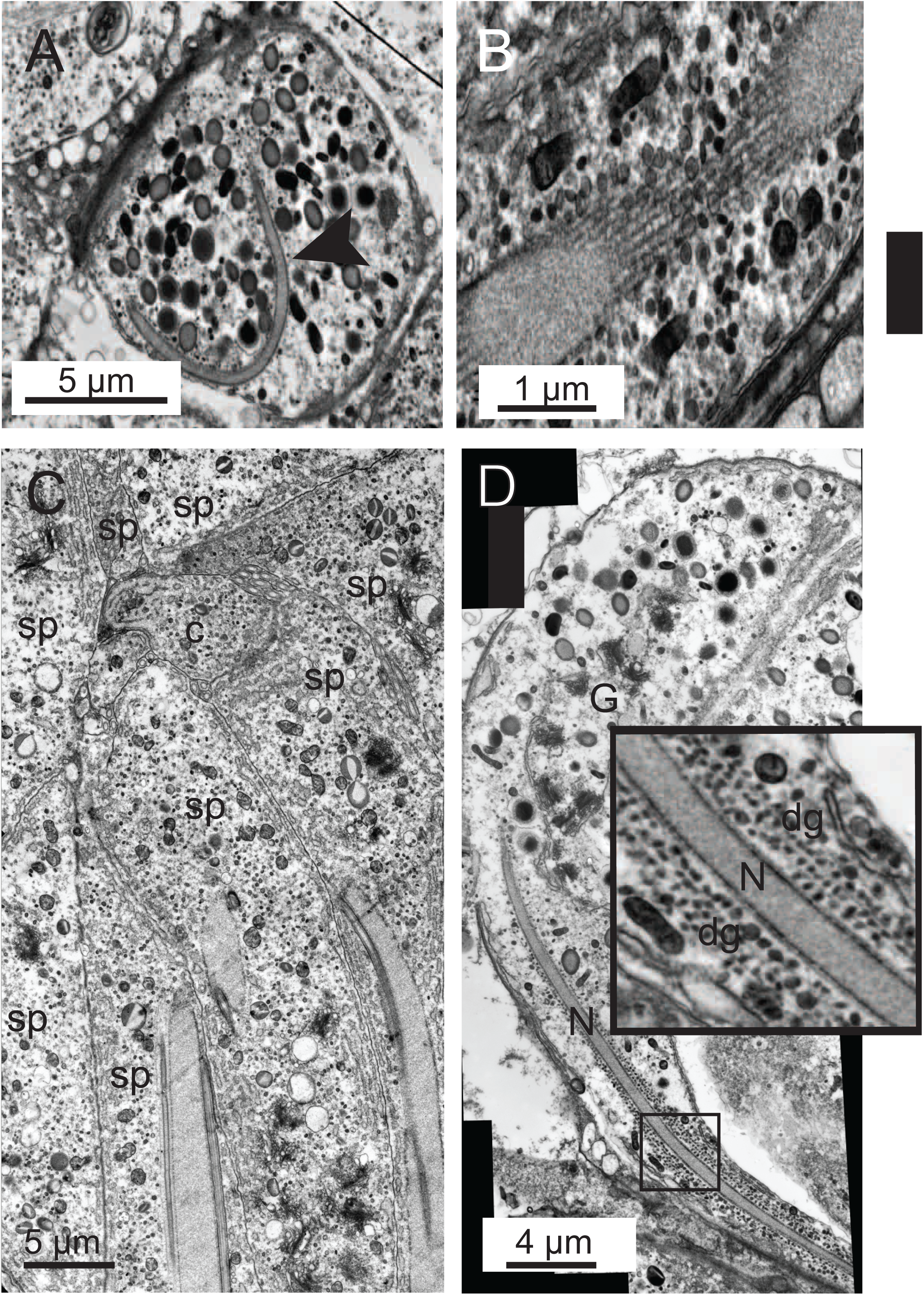
Early stages of spermiogenesis. A: A primary spermatocyte containing an elongated, curved nucleus (black arrow). The cytoplasm is full of large granules. B: A close-up of a similar spermatocyte nucleus cut twice as it passes through the section. The manchette of single microtubules is clearly visible surrounding the nucleus. C: A montage image showing a later stage in spermiogenesis. Several spermatocytes (sp) are seen arranged around a common cytophore (c). The nuclei are elongated and the partially internalised flagella are intimately associated with them. D: A different orientation showing the cytophore containing large numbers of Golgi (G) and developing large granules and an elongating spermatocyte. At this stage the manchette of microtubules surrounding the nucleus (n) becomes associated with huge numbers of dense granules (dg) (see inset).

In cross-section, the early-stage spermatocytes contain an ovoid nucleus, but in more mature testes they appear elongated and become crescent shaped (Fig 3A), each surrounded by a complex *manchette* of single microtubules that enclose the nucleus in a cage-like structure (Fig1-C, Fig 3B). The nuclear envelope presents as a fragmented series of individual cisterns separated by gaps. Each of these cisterns is associated with a single manchette microtubule. Approximately 50% of the nuclear material appears to be exposed to the cytosol (see also Fig A-E and Fig5B). We did not see many examples of centrioles in the cytoplasm of early stage spermatocytes, but it has been suggested from related species that they migrate from near the plasma membrane towards the nucleus [34].

The spermatocyte extends and the nucleus elongates. The flagella are simultaneously incorporated into the cytosol of the spermatocyte progressing in a ‘disto-proximal’ mode (Fig 1 D-E, Fig 3C).

The resultant spermatozoon has a long, thread-like (filiform) nucleus which, by moving distally as the spermatid matures, but not passing between the centrioles of the developing tail, comes to lie in front of the incorporated binary flagella (Fig 2).

Sections through the elongating spermatocyte at different stages reveal that both flagellar axonemes come to be associated with the elongating nucleus. Flagellar incorporation is asymmetrical. At early stages in spermiogenesis one axoneme is predominantly ’free’ in the cytosol whilst the other is entirely surrounded by a cytoplasmic canal or ’sheath’, a thin <20 nm luminal void within the cytoplasm generated as the flagellum is drawn into the cytoplasm (Fig 4 A). The internalised flagellar sheath appears to be held close to the nucleus by association with a single microtubule (Fig 4 A-C) which runs between it and the nuclear envelope. The cytoplasmic ’free’ axoneme is also associated with the nucleus but in this case by a ring of approximately 12 satellite single microtubules which surround it (Fig 4 D-E).

**Figure 4.**
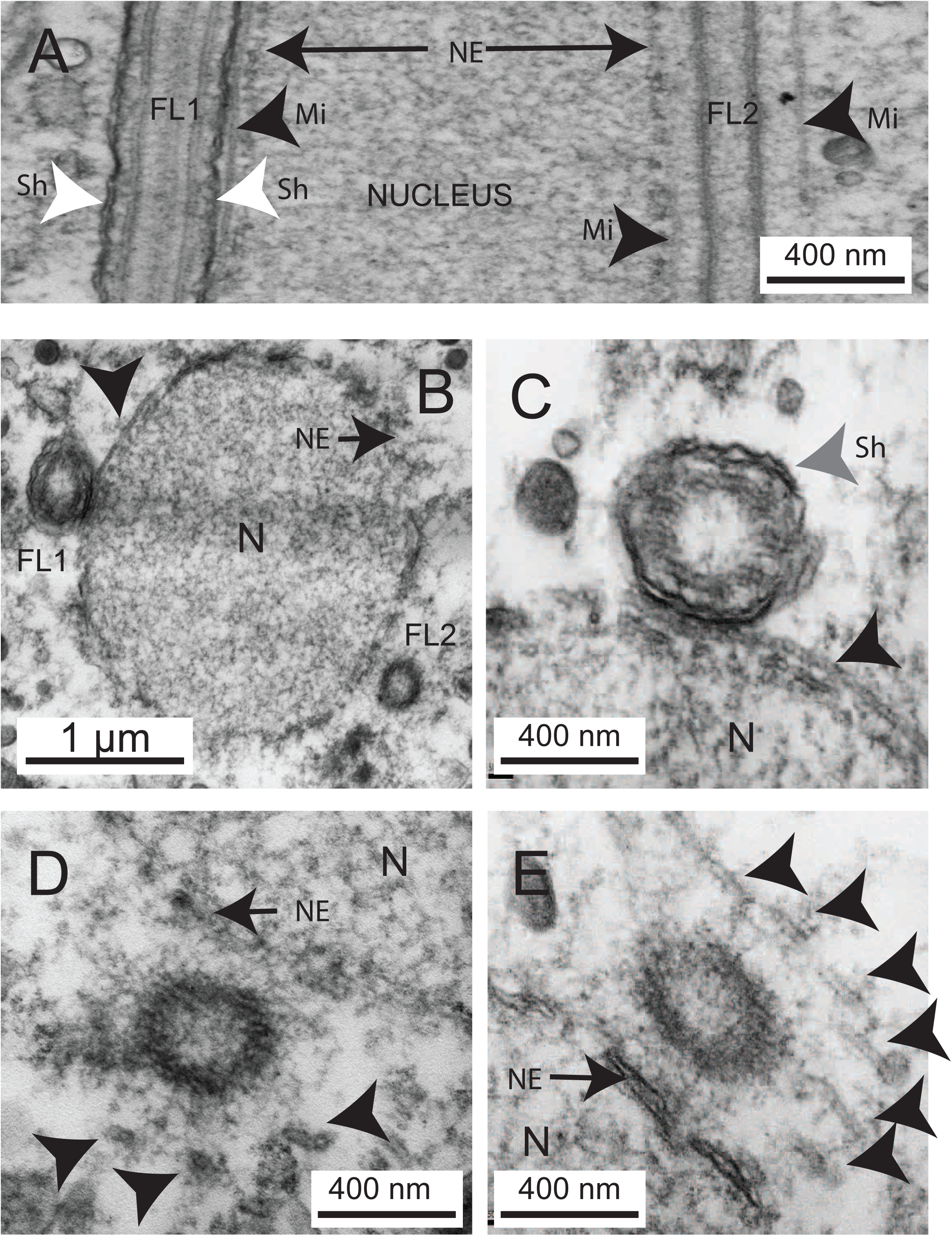
Flagella axoneme association with the nucleus. A: longitudinal section through a primary spermatocyte. Internalised flagellum to the left (FL1) showing the sheath of plasma membranes (Sh and white arrow). A single microtubule (Mi and black arrow) lies in the space between the flagellum and the nuclear envelope (NE). The non-internalised flagellum (FL2) to the right is associated with the nucleus via with a ring of microtubules of which two are observable in this section here (Mi black arrows). B: Transverse section of a different primary spermatocyte showing the same features identified above. C: A close-up from another specimen showing the dual membranes of the cytoplasmic canal/sheath (Sh) grey arrow of the internalised flagellum. The nucleus (N) is surrounded by a highly porous nuclear envelope. D and E: Non-internalised flagellum associated with the nuclear envelope (NE) of the nucleus (N) by a ring of microtubules (black arrows).

As spermiogenesis proceeds, the remaining external projections of both flagella are drawn into flagellar sheaths. These internalised extracellular domains progressively reduce in volume to form narrow canals that no-longer encircle the entire flagellum. The result is that ultimately both the flagella axonemes are effectively ’free’ in the cytosol and flagellar incorporation is complete. The flagella migrate past the nucleus and are fully incorporated as lateral axonemes, 9 + 0 rings of microtubule doublets on opposite sides of the sperm (Fig 5A). They are no-longer associated with the nucleus, which has moved beyond them.

**Figure 5.**
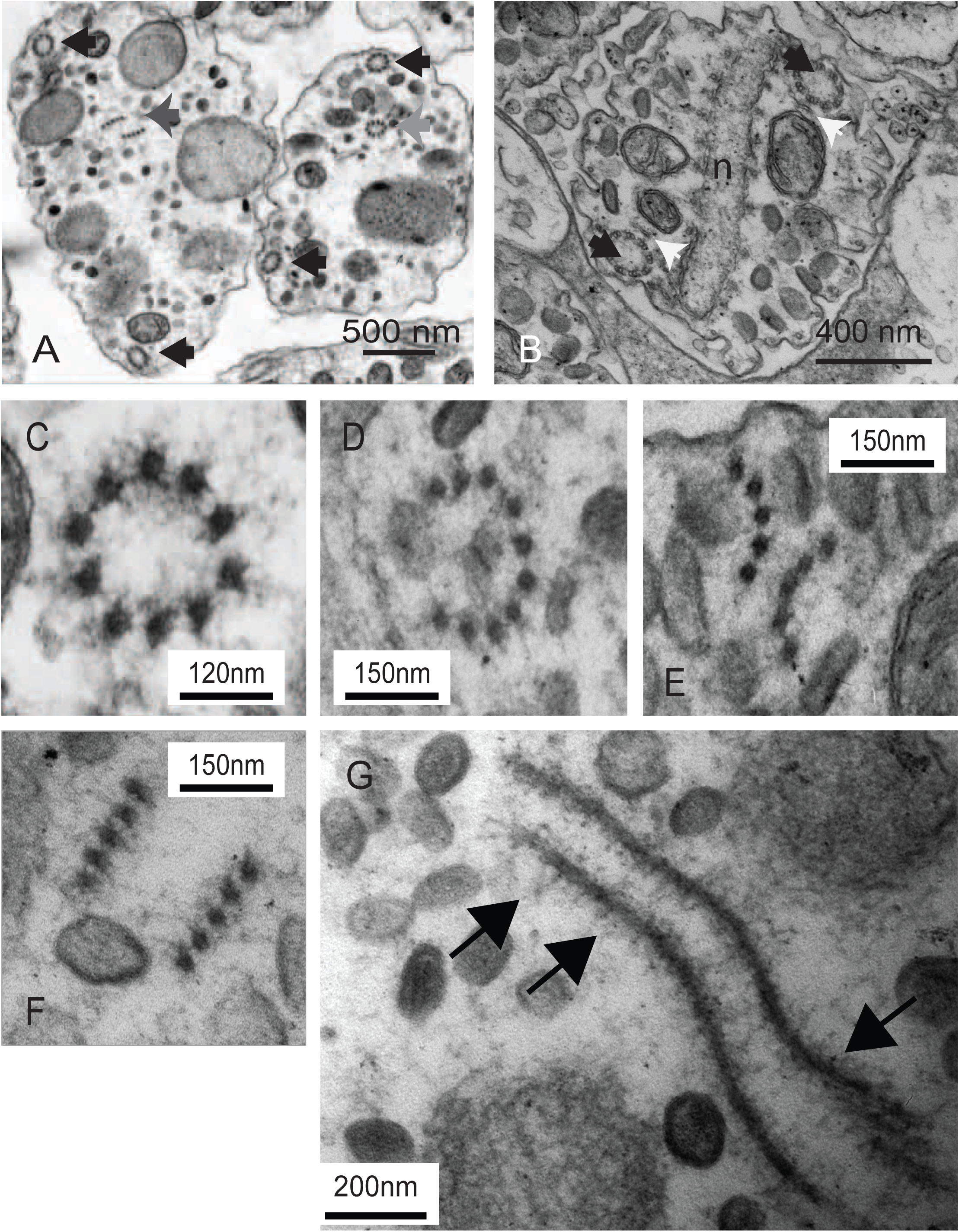
Arrangement of the medial axoneme (keel) at different stages of development. A: Later in spermiogenesis the flagella (black arrows) lie peripherally in the mid-portion of the spermatocyte. The internal axial keel (grey arrows) is located nearer the centre. B: The nucleus (seen here in a more mature spermatocyte) is flattened. The flagellar axonemes are fully internalised (black arrows) and the flagellar sheath is much reduced (white arrows). C: The medial axoneme or *keel* is a tubular array of 10 atypical microtubules. D: More distally the ring breaks and becomes an open crescent. E: This, in turn, breaks down into two sets of 5, initially arranged as two arcs. F: The rods become aligned as two staves. G: Staves seen in transverse section. Note the dense array of radiating filaments (black arrows). Note the filamentous extensions on the staves which remain even though the dense granules are no longer associated with them.

The nucleus elongates becoming flattened and condensed (Fig 2 and Fig. 3D). Down the centre of the spermatozoon is a so-called ‘medial keel’, a ring of 10 microtubule-like rods. These do not show the typical doublet ring of microtubules of conventional axonemes or appear as single microtubules. They are larger than the latter and have no internal cavity. We refer to these as dense core atypical microtubules. Over the course of spermiogenesis this medial keel reorganises, firstly into two semi-circular bundles of 5 and ultimately into two linear ‘staves’ (Fig 5C-G, Fig. 7A-B).

**Figure 6.**
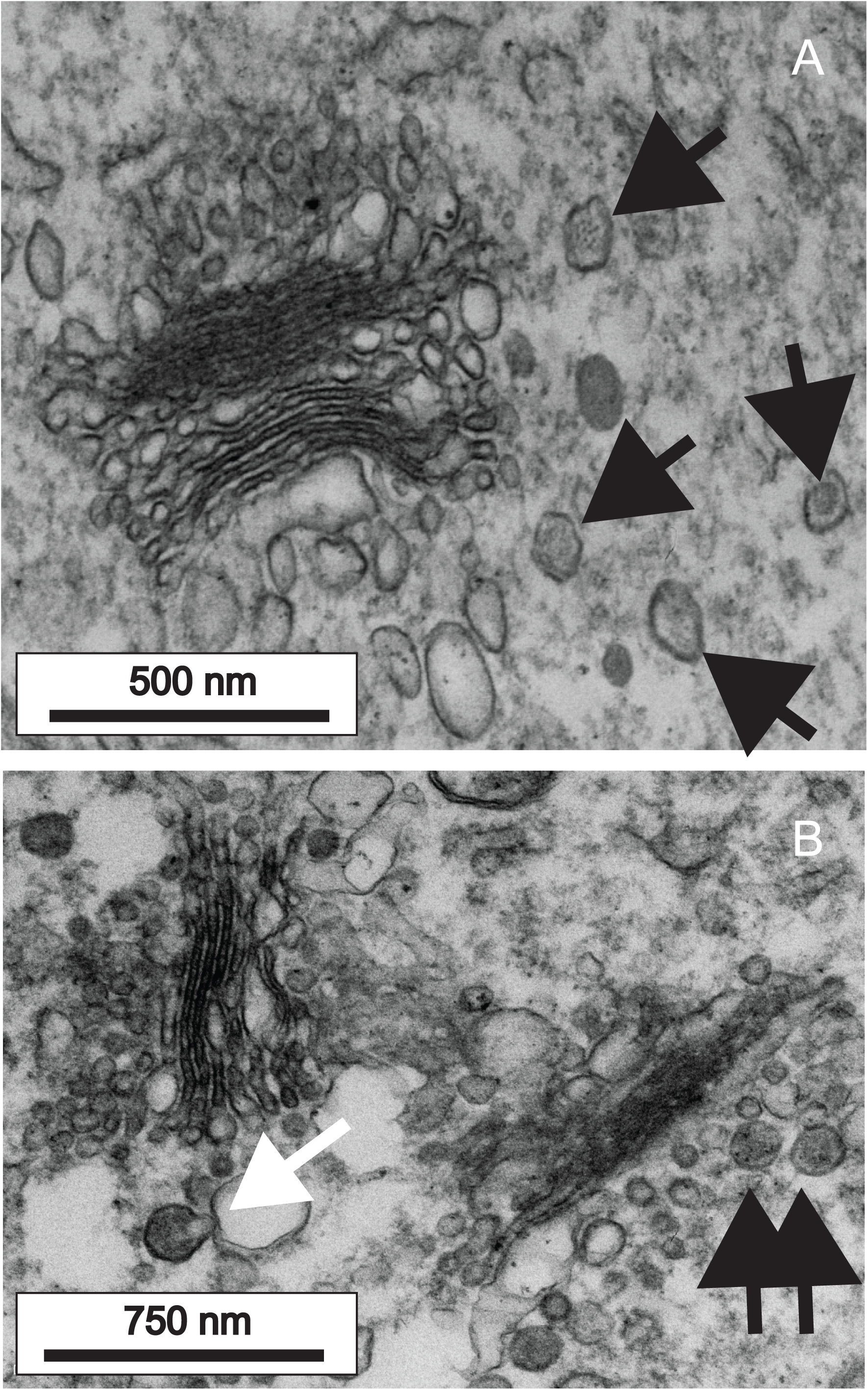
Dense granules are derived from the Golgi. A: Golgi apparatus in an early spermatid. Note dense granules containing filaments in the vicinity of the trans Golgi network (black arrows). B: Two Golgi apparatuses in an early spermatid. Fully-formed dense granules are shown (black arrows) as well as a dense granule apparently budding from a large vacuole of the trans Golgi network (white arrow).

**Figure 7.**
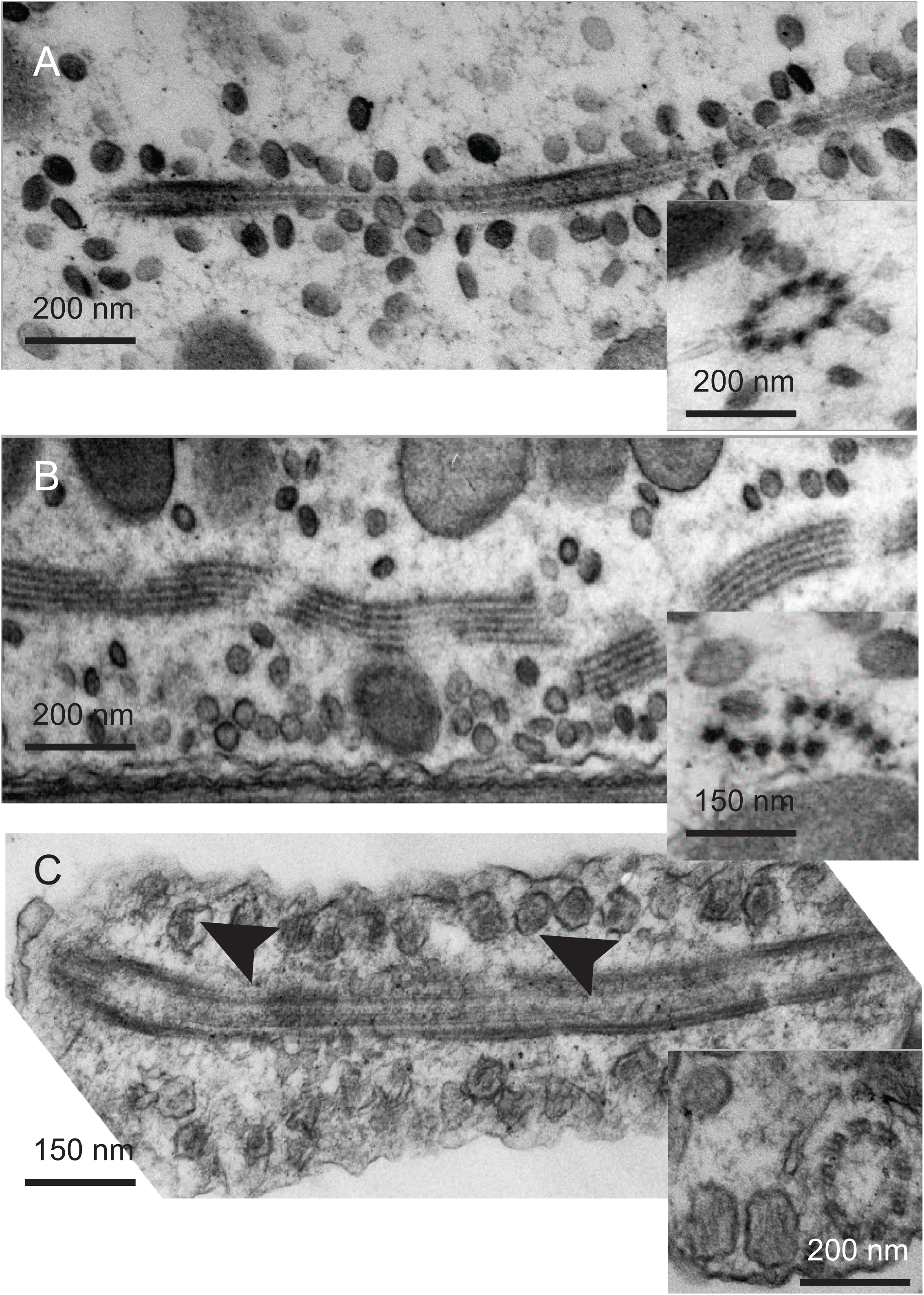
Movement of dense granules from the medial to the cortical axonemes. A: In immature spermatids the (DGs) are arranged spirally (see inset) around the circularly bundled, axial axoneme. B: Later in development the axoneme unfolds into two linear structures (a microtubular stave: see also inset). The DGs leave the axoneme and become associated with the cortical plasma membrane. C: In the vicinity of the cortical axoneme the DGs (black arrows) become aligned to the plasma membrane, their electron-dense contents perpendicular to the membrane surface.

### Nuclear envelope-plasma membrane contact: formation and trafficking of dense granules

Large numbers of small (50nm) dense granules (DGs) were observed surrounding Golgi apparatuses in primary cytophores and primary spermatocytes (Fig 6). These DGs have a characteristic structure, being flattened discs with a thickened circumferential rim. In the immediate vicinity of the Golgi we observed vesicles showing internal structures of varying degrees of condensation; some appearing as punctae, others as fibres and yet others as dense aggregates. We interpret these as immature DGs in the process of condensation, their contents, upon aggregation or polymerisation, ultimately imposing structure on the mature discs. These granules ultimately contribute to the nucleus plasma-membrane contact site.

The mature DGs associate with microtubules of the manchette of the nucleus (Fig 3D) and also form a spiral array around the circularly-arranged medial keel axoneme (Fig 7A, Fig 1-F), to which they are attached by fibrous tethers (not visible in all sections). It is possible that the medial keel axoneme is continuous with the nuclear manchette microtubules at this stage (as indicated schematically in Fig 1 and Fig 2). We do not know if DGs are transported down this structure or merely stored here at this stage of spermatogenesis. Similar granules were previously identified in a related species [12]. At this stage of spermatozoon development there are very few DGs in the cell periphery.

During the process of axial keel remodelling into two staves (Fig 5C-G, Fig 7A-B) (described later), the DGs dissociate from them, becoming re-distributed adjacent to the plasma-membrane in the immediate vicinity of the lateral axonemes (Fig 7C).

### Localisation and formation of the nucleus-plasma membrane contact site: dense granule maturation

As spermiogenesis proceeds approximately 50% of the dense granules appear to be in a state of abeyant fusion with the plasma membrane (Fig 7C and Fig 8A). Of these 71% (n=200) were clearly aligned with their internal filaments arranged perpendicular to the plasma membrane. Of the remaining ∼25% it was not always possible to definitively establish the orientation of the plasma membrane. The majority of these dense granules are in the immediate vicinity of the laterally internalised flagellar axonemes (94.7%, n=200); or on the residual cytoplasmic canal that separated these axonemes from the cell body (3.5%). There were very few dense granules associated with plasma membrane away from these two sites (1.8%). Despite the finding of dense granules along the plasma membrane, the fibrillary material that is found around the nucleus is never found associated with the plasma membrane unless the nuclear envelope is contacting it. suggesting that the dense granules only deliver their contents when contacted by the nuclear envelope; contact with plasma membrane alone is insufficient to elicit its release.

**Figure 8.**
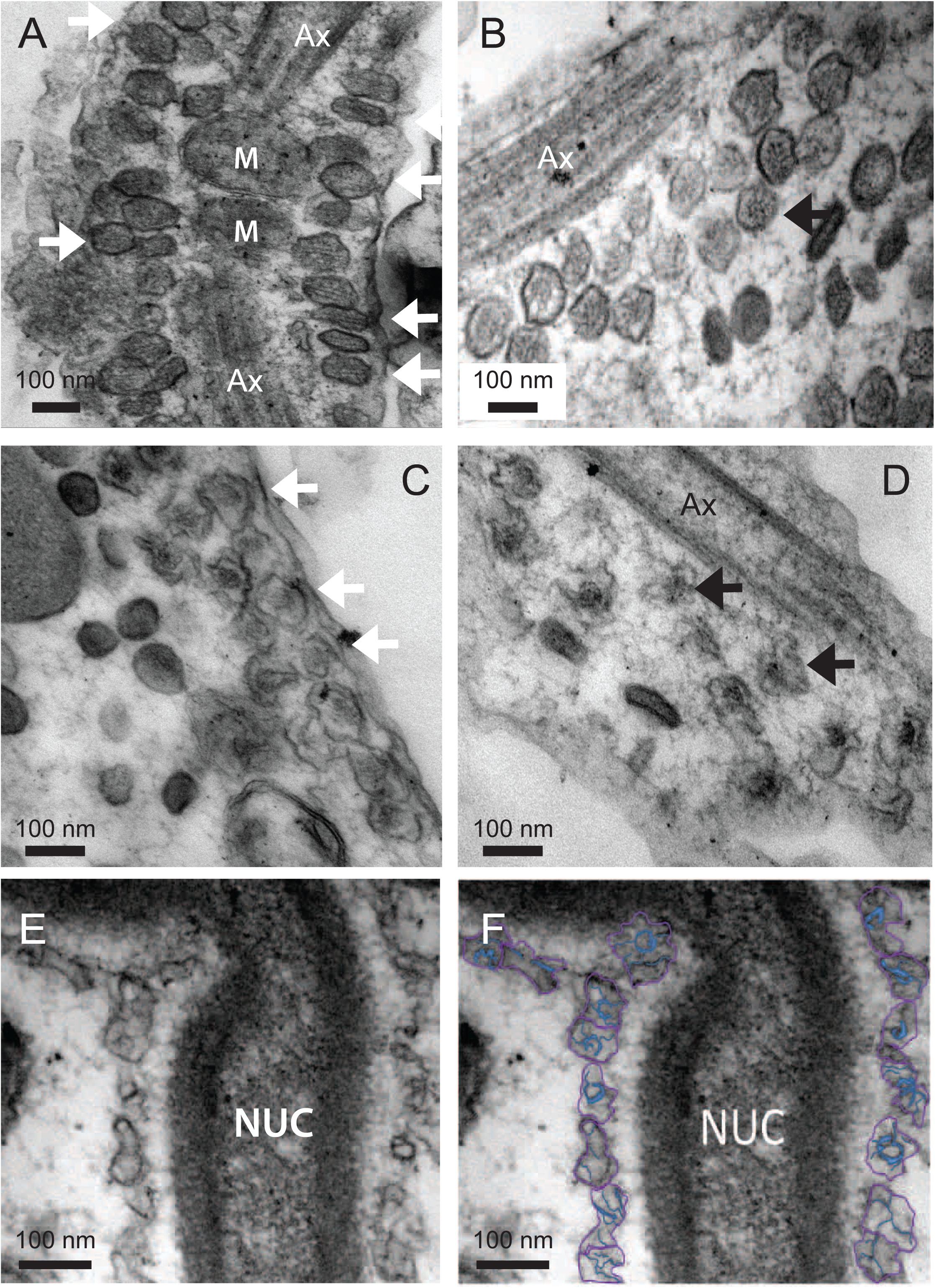
Conformational changes in the dense granules (DGs) A: DGs aligned with the plasma membrane in the vicinity of the cortical axonemes (Ax). Mitochondria are also associated with these structures (M). B: The surface of the DGs (white arrows) deform, flatten and become crenelated and some of the filaments shorten. C: Flattened granules seen obliquely. Most electron dense filaments collapse to a central core whilst others remain attached to the DG membrane (white arrow). D: The DGs continue to flatten (note the vicinity to the axoneme). E and F: DGs become associated with/incorporated into the nuclear envelope (DG membranes here outlined in purple, the internal filaments are high-lighted in blue).

The dense granules become incorporated into the peri-nuclear layer (Fig. 8E-F and Fig 13). Here we look at some detail of the contents and shapes of dense granules.

Dense granules associated with the plasma membrane have a narrow neck region when viewed from the side and a dense circular connection to the plasma membrane. Between 10 and 18 filaments, around 3 nm in diameter are packed hexagonally, emanating from a point close to the plasma membrane then running parallel before coming together again. (Fig 8A and Fig 9). This structure seems to be cross-linked by fine lateral filaments. The dense granule membrane, initially circular in outline (Fig 8A-B), becomes progressively more undulating as spermatid maturation progresses (Fig 8C-D and Fig 9A-F) and the 3 nm internal filaments crowd together in the centre. Later still, the filaments condense further, becoming shorter and the membrane expands to become flower-shaped (Fig 8D and Fig 9E-F).

**Figure 9.**
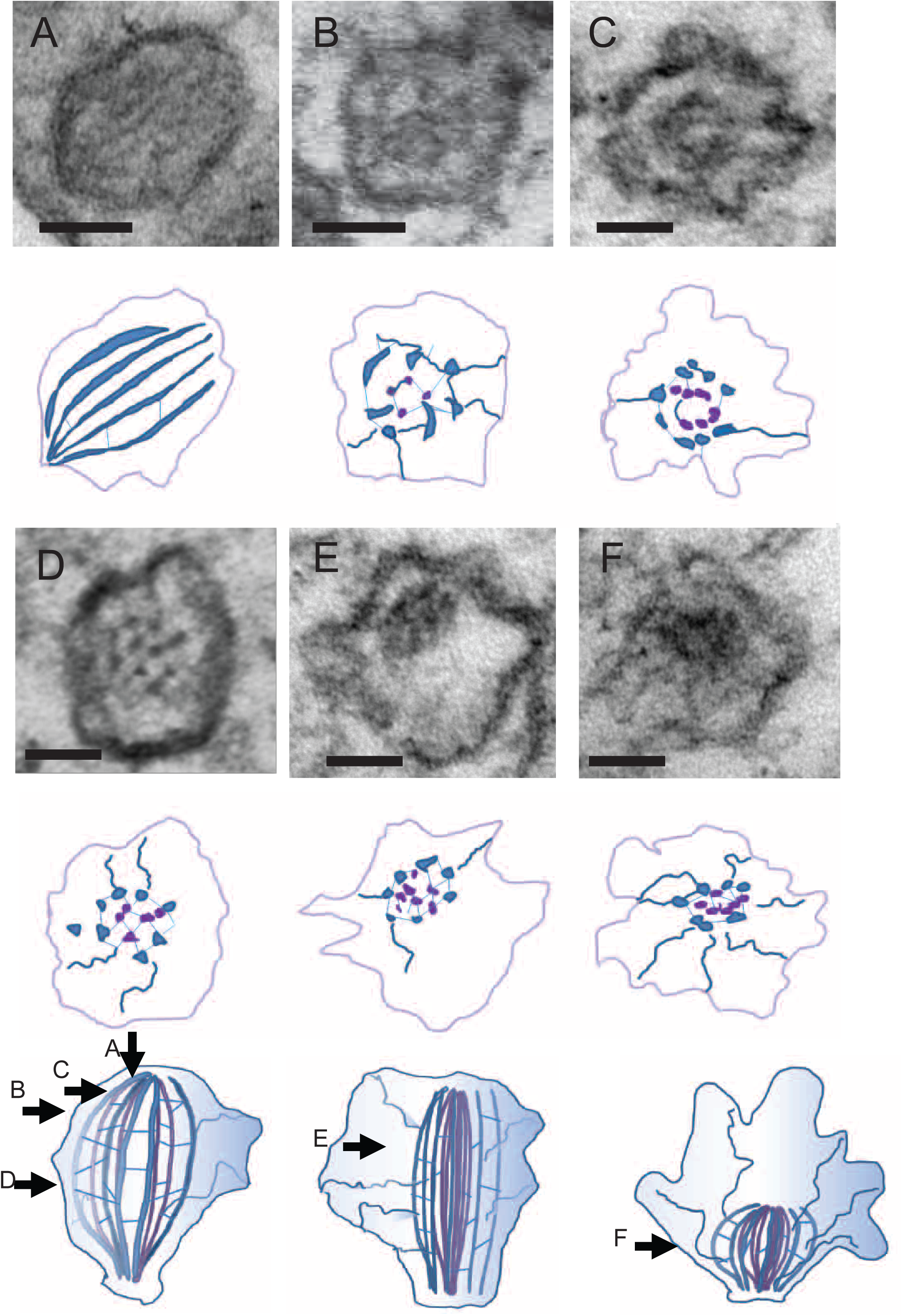
Schematic: demonstrating the condensation of the dense granules into the multicomponent nucleus-plasma membrane contact site. A: A dense granule aligned with the plasma membrane observed perpendicular to the plasma membrane showing the internal filaments (blue). B: A granule at a similar stage of fusion sectioned transversely showing arrangement of an additional layer of internal filaments (purple) inside the outer layer (blue). Radiating filaments which reach out to the membrane are also clearly seen. C: A slightly later stage: the outer membrane is crenelated and internal filaments shorten and aggregate. D: A granule cut transversely at an early stage of fusion showing the internal filaments equidistantly spaced. Very thin horizontal connections can be seen interconnecting the filaments. Presumably these retain the structure’s geometry. E: Later the filaments coalesce and are bundled asymmetrically in the granule. F: A late-stage granule cut transversely. The internal filaments condense to dense puncta. In this example the radiating fine filaments are quite prominent. The schematic at the bottom of the figure is a graphic representation of where these sections have been taken.

As spermatogenesis progresses, the nuclear envelope of the spermatozoon, which was characterised by isolated cisternae separated by large gaps where the nuclear content was exposed to the cytoplasm, becomes more elaborate, and florid extensions of the membrane develop. These proliferations hold the long, filiform nucleus suspended through the centre of the sperm cell by connections to the two opposing faces of the plasma membrane (Fig 10A). This led us to wonder where the additional membrane came from and how the nucleus was able to form an attachment with the plasma membrane at these sites. On close examination it appears that the nucleus forms a contact site with the dense granules, possibly fusing and incorporating their membrane into that of the nuclear envelope ruffles. The DG membrane is slightly more osmophilic (appearing more electron dense) than that of other regions of the nuclear envelope, and in regions where fusion seems to be actively occurring, remnants of DGs are visible (Fig 8-E-F). In many sections we could identify intermediates in the process of fusion and we also observe rings of electron-dense particles we interpret as the remnants of the DG dense fibres (Fig 11 and insets). This process was easier to observe in samples processed to enhance membrane staining (Fig 12). This entire process is summarised schematically in figure 13.

**Figure 10.**
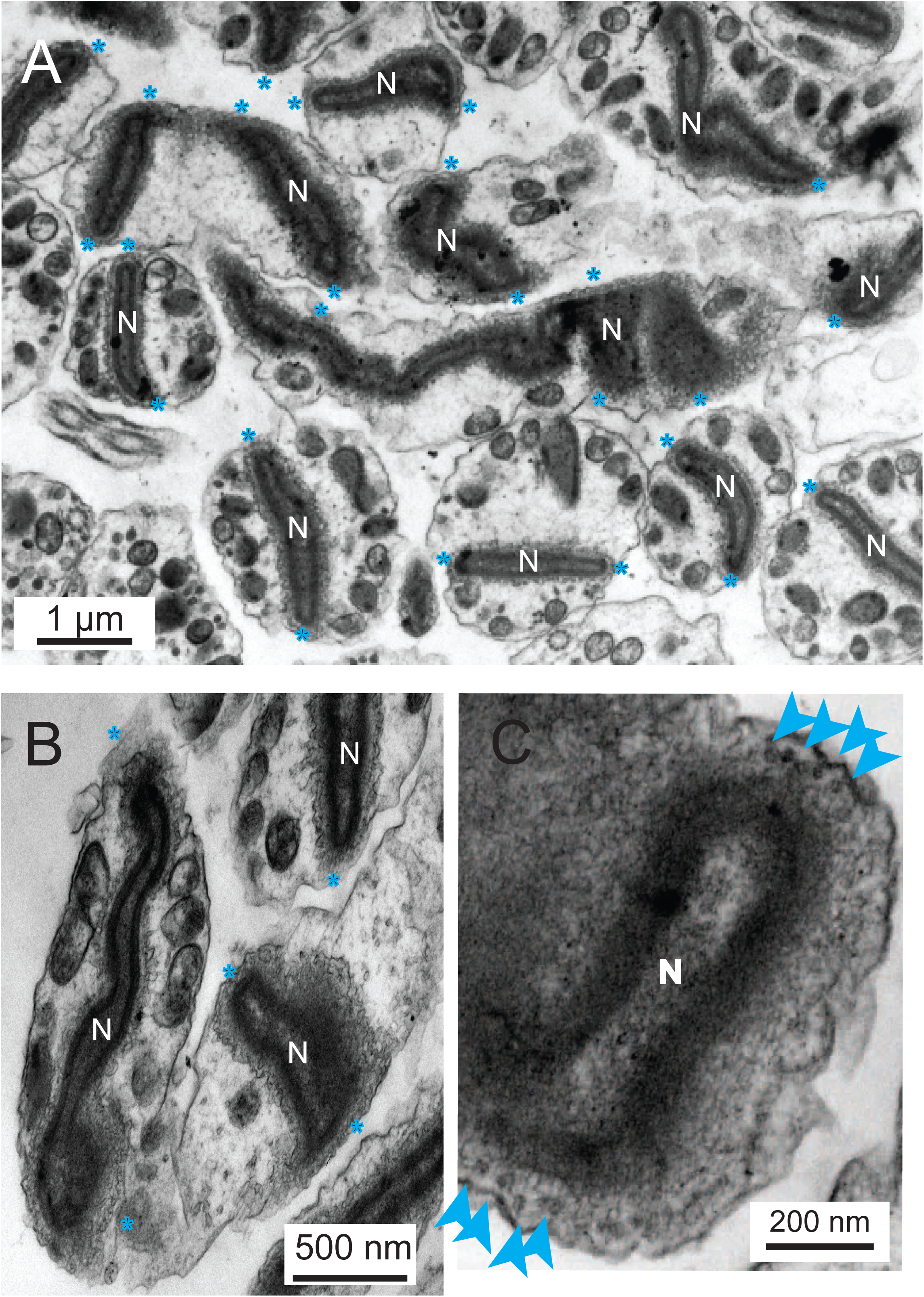
The nuclear envelope retains elements of the fusion events. A: In the mature spermatozoan the nucleus (N) appears ‘suspended’ in the middle of the cytoplasm as a result of a number of contacts with the plasma membrane (blue stars (*)). B: More examples showing the complex elaborations of the membranes. C: Contact sites are characterised by (20nm) rings of electron dense puncta (blue arrows).

**Figure 11.**
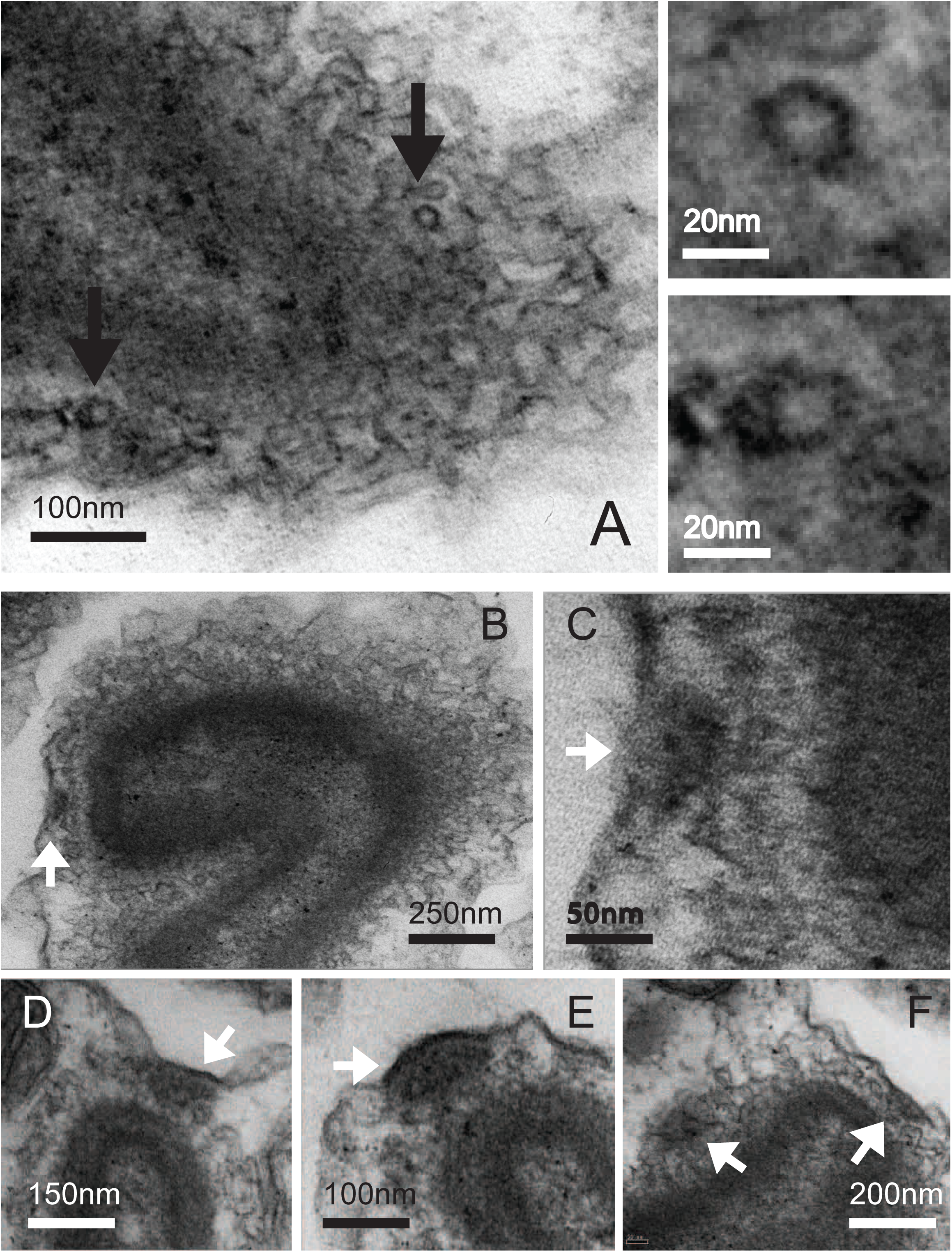
A: The (20nm) rings of electron-dense puncta (black arrows) and insets. B-F: Rarely we also observe additional aggregate densities which also appear to link the nucleus to the plasma membrane (white arrows).

**Figure 12.**
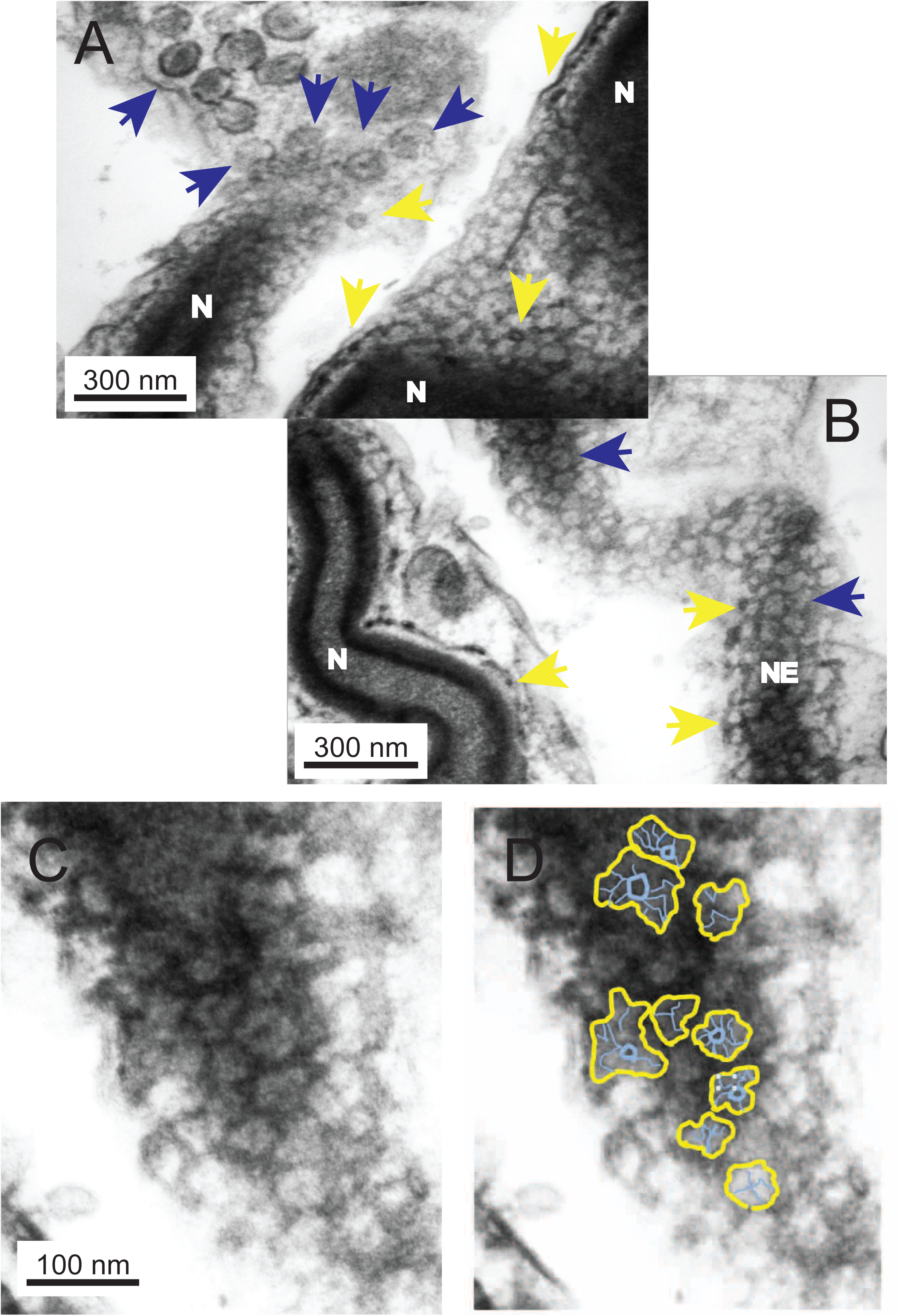
A preparation to enhance membrane staining facilitates contact site interpretation. A-B: A mature spermatozoan, sectioned to provide a glancing slice through the nuclear envelope. The unincorporated dense granules can be seen in the vicinity of the encroaching nucleus (blue arrows). Incorporated DGs are reduced to rings of electron dense puncta (yellow arrows). C-D (schematic): The filaments (blue) and membranes (yellow) of the DGs are incorporated into the nuclear envelope plasma-membrane contact site. N = nucleus, NE = nuclear envelope.

Less frequently, there were larger, more complex, focal accumulations of material within nucleus-plasma membrane contacts. These varied in number between specimens, so we conclude they represent a transitory state in spermiogenesis (Fig 11 B-F).

### Flagellar incorporation is associated with formation of another nucleus-plasma membrane contact site

At the point of entry of the internalised flagella into the developing spermatocyte, a multi-laminate, electron-dense complex (MEC) forms between the flagellar sheath and the nucleus (Fig 14A). This structure maintains an intimate association of the flagella with the nucleus as nuclear elongation and flagellar incorporation proceeds (Fig 14B-E, schematic Fig 1D-E). The juncture of each internalised flagellum and the nucleus is marked by an electron-dense membrane contact site (Fig 14D) where the nuclear envelope appears to be fixed to the MEC and associated internalised plasma membrane.

**Figure 13.**
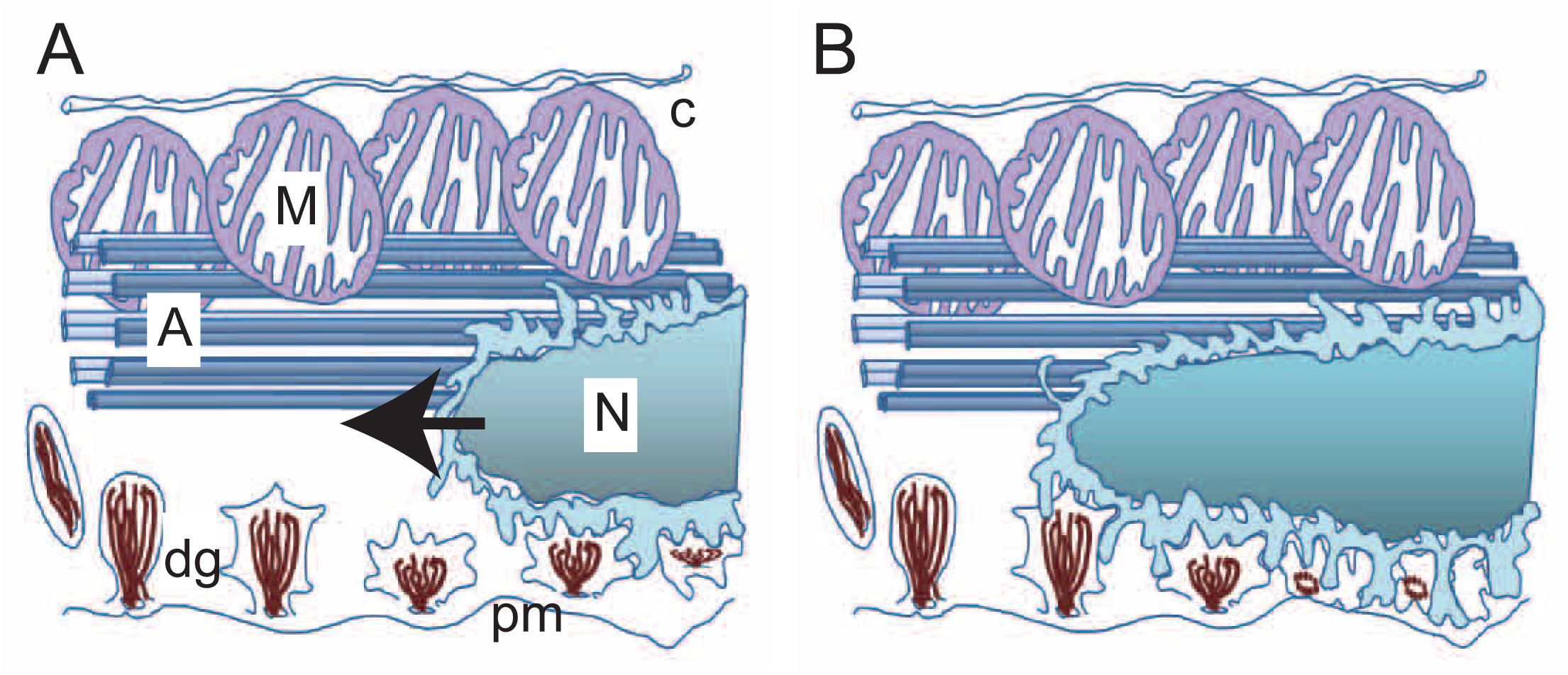
Proposed model of nuclear migration leading to incorporation of dense granules into the elaborating nuclear envelope (N = nucleus, A = axoneme, M = mitochondrion, dg = dense granule, pm = plasma membrane).

**Figure 14.**
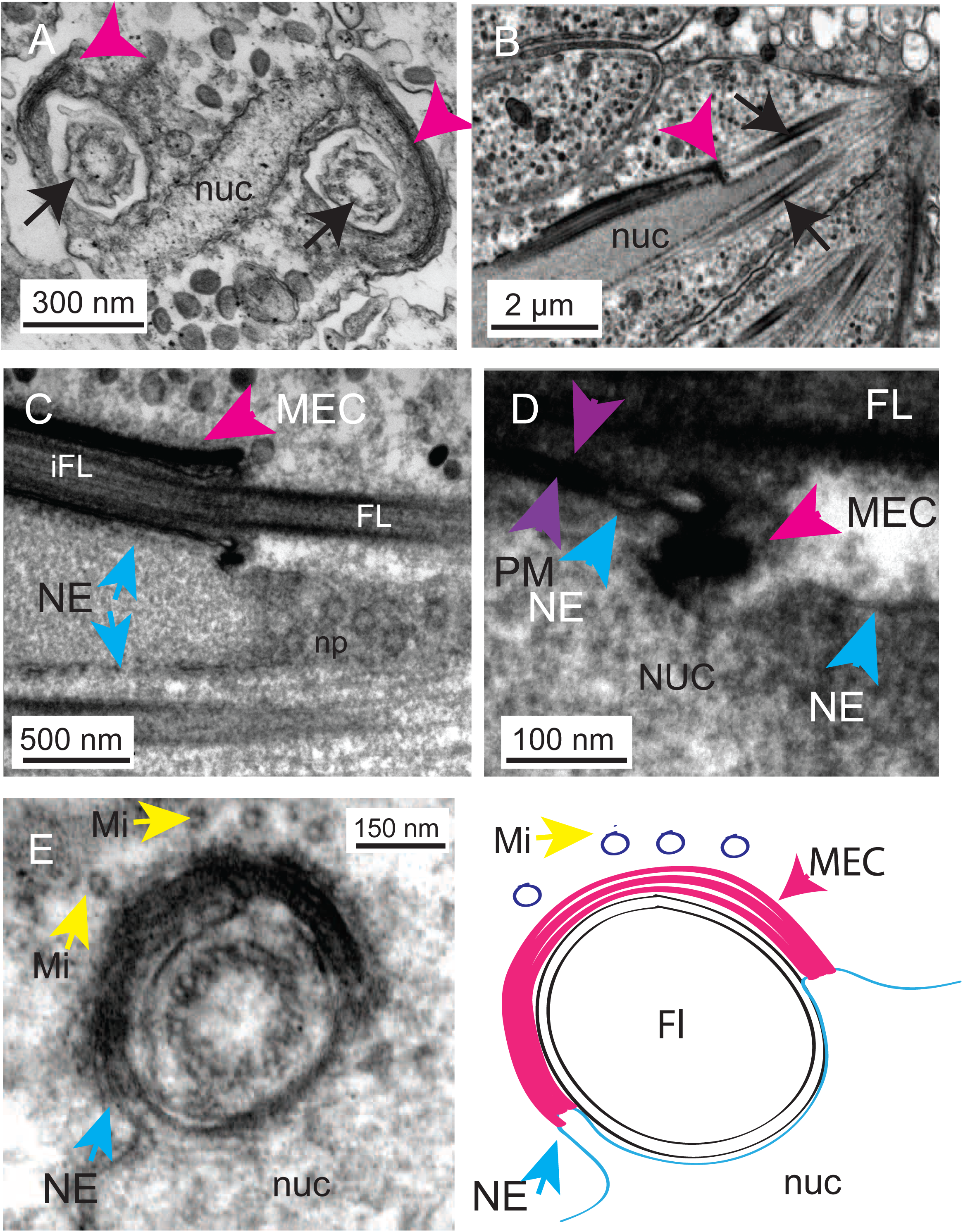
A multilaminate, electron-dense complex (MEC) (a nucleus-plasma membrane contact site). A: A transverse section of the tip of the developing spermatocyte showing the flattened nucleus (nuc) and development of a multilaminate, electron-dense complex (MEC) (pink arrowheads) that forms as the flagella (black arrows) are drawn into the developing flagellar sheath. B: A transverse section showing the internalising flagella (black arrows) and the association of the MEC (pink arrowhead) with the nuclear envelope (blue arrows). C: The MEC is intimately associated with the nuclear envelope (note the nuclear pores (np) seen in this glancing section across the nucleus). D: A close up of this region showing the contact site between the sheath (composed of invaginated plasma membrane (purple arrowheads) and the nuclear envelope (blue arrowheads)). E: Note the ‘drawing in’ of the nuclear envelope (NE and blue arrow) into the MEC. There are microtubules associated with the MEC complex at this stage (Mi yellow arrows). E: Schematic to show the arrangement of membranes at the nuclear envelope/sheath contact site mediated by the MEC. nuc = nucleus, NE = nuclear envelope, Mi = microtubule, FL = flagellum, iFL = internalised flagellum, MEC = multilamellar electron-dense complex.

An MEC has also been observed in electron micrographs of spermatocytes of *Symsagittifera bifoveolata* and *Symsagittifera schulzei* [24]. This is presumably a temporary structure as it is not seen later in spermatogenesis.

### The medial keel is a centriolar derivative and links the apical nucleus to the tail and to the lateral axonemes

Examination of large numbers of sections of mature sperm revealed a coordinated pattern of microtubule reorganisation passing spatially from the nucleus to the tip of the tail. In the main body of the sperm the lateral axonemes are equatorial on either side; but towards the tip, as the tail narrows, they come to lie closer together until there is a clear dorsal/ventral loss of symmetry. The flagella appear as two 9 + 0 axonemes of typical microtubule doublets side by side. In this region, the dense-cored atypical microtubules of the medial keel axoneme separate and are reduced to smaller bundles distributed equally dorsally and ventrally. Towards the very tip of the sperm there is a progressive drop-out of microtubules until only a couple remain above and below the paired axonemes (Fig 15A i-ii). One of the lateral axonemes progressively loses its 9 + 0 doublet structure and appears as a 9 + 0 ring of atypical microtubules identical in structure to the remaining axial keel solid-cored microtubules. Distal to this point, this axoneme ring of dense-cored atypical microtubules loses its circularity and progressively loses tubules (Fig 15Aiii-vi); we see the appearance of a new atypical microtubule in the centre of the remaining intact lateral axoneme (which is still composed of conventional doublet microtubules), and yet more distally, this is joined by a second internal tubule (Fig 15A vii-viii). Given that we rarely saw such sections, it is likely that this is a relatively short domain and, being composed of a so-called 9+”2” arrangement of microtubules (a ring of 9 normal doublets plus two atypical microtubules in the core), it is likely to be the motile region of the sperm tail, the encircling doublets being able to derive force by interaction against the internal core. Beyond the proposed motile section, the doublets of the remaining lateral axoneme also assume the form of atypical microtubules, these subsequently being lost until all that remain are a swirl of tubules composed, we suspect, of both axial keel and lateral atypical tubules (Fig 15A ix-x).

**Figure 15.**
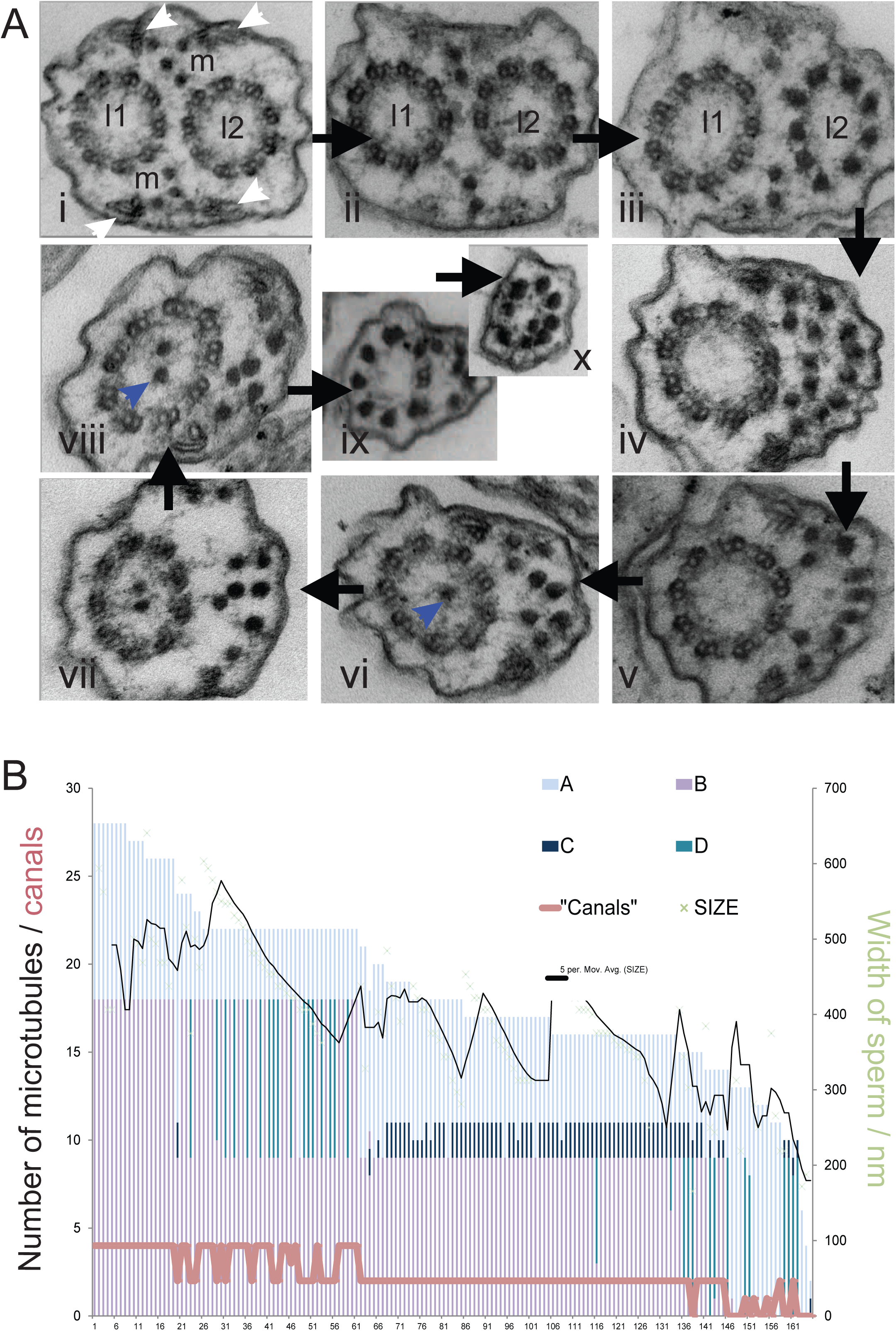
Remodelling of cortical axonemes is coincident with their conversion into microtubule staves. Ai-ii: The spermatozoan tail loses microtubules structures towards the tip. At its thickest, the two lateral, cortical 9 + 0 flagellar axonemes align side by side (l1 and l2). The axial keel axoneme (initially 2 pairs of 5 staves) is reduced from 5 then 4 atypical dense-cored microtubules which align apically and basally to the lateral axonemes (m).There are also 4 other membranous “canals” associated with microtubules. These are probably the remnants of the flagellar sheath (white arrowheads). Progressing towards the tip of the sperm tail the keel microtubules reduce in number until a critical point at which one of the lateral axonemes converts into atypical dense-cored microtubules (iii). The circular arrangement is lost and they intermingle with the microtubules of the axial axoneme (iv-vi). At this point a new microtubule develops in the centre of the remaining cortical axoneme (blue arrows) (vi) this is joined by another to provide a 9 + 2* axoneme. This axoneme too converts into the dense cored microtubular form. Microtubules are lost towards the tip of the spermatozoon. B: Histogram showing the number of microtubules (doublets counted as two, atypical microtubules as one) in many serial sections of spermatozoon tail. The green line represents the thickness of the tail at each point. Keel microtubules are coloured blue (A), lateral axoneme doublet microtubules are coloured purple (B), internal atypical microtubules are dark blue/black (C), lateral, atypical microtubules are blue-green (D). The number of “canals” is shown in pink. Progressive loss of microtubules occurs as the tail becomes thinner; but there are specific regions where doublet microtubules convert to atypical ones.

Loss of microtubules correlates with the decreasing thickness of the tail and there are distinct regions where microtubules tend to convert from one form to another. Fig 15B.

We were able to identify intermediate microtubule structures between the canonical microtubule ‘doublets’ and the atypical, dense-cored microtubules (Fig 16). In some cases these appear as doublets with additional extensions or triplet rings, leading to the possibility that the dense-cored microtubules could be derived by rearrangement of triplet structures. These successive changes in axonemes over distance suggest that multiple changes in microtubule structure and subsequent bundling arrangements may arise from simple gradients of one or more components.

**Figure 16.**
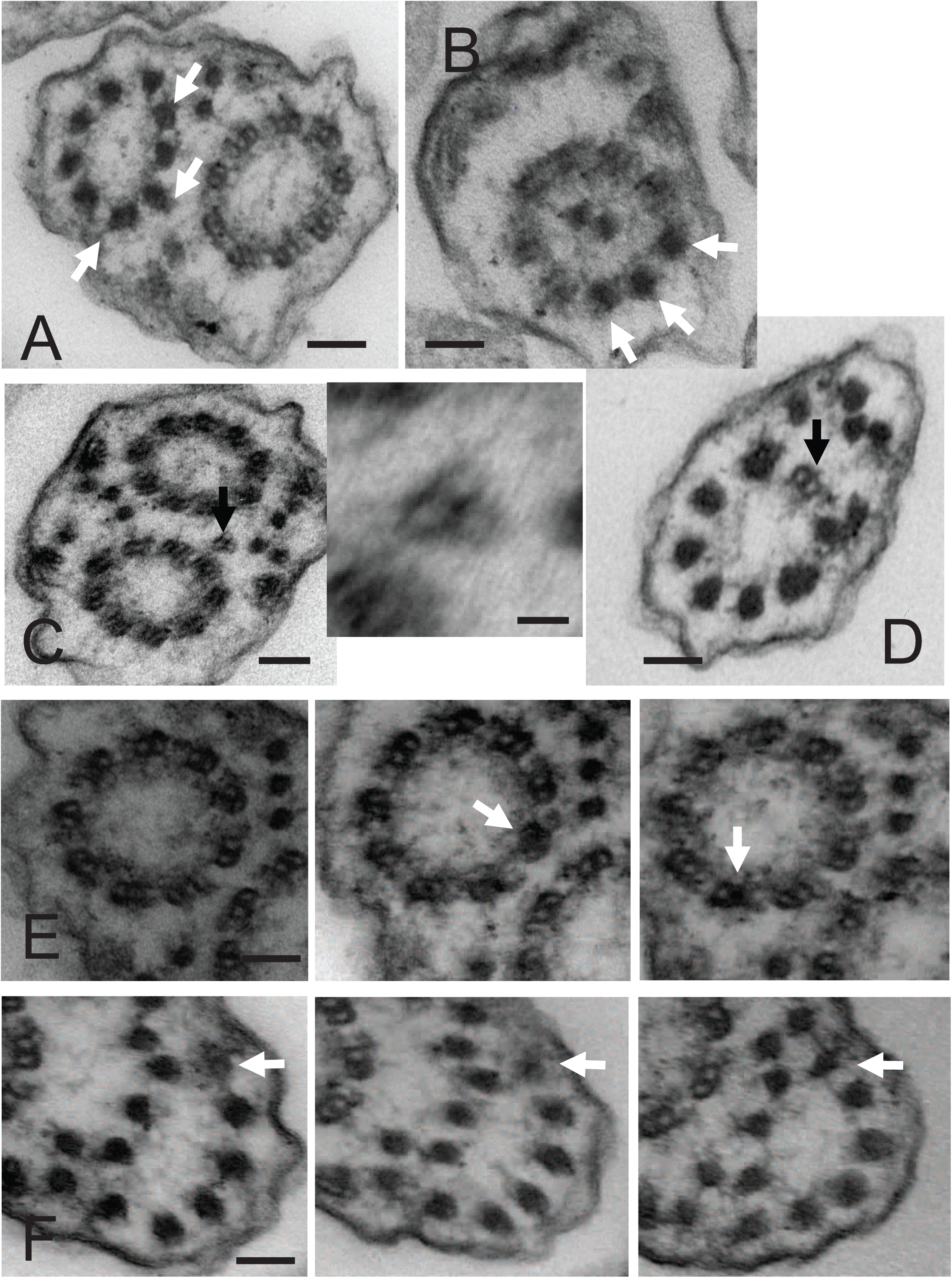
Interconversion of microtubule doublets to atypical dense core microtubules. A: A transverse section of a spermatozoon tail showing one of the cortical axonemes in which three of the microtubule doublets show characteristics intermediate with atypical dense-core microtubules (white arrows). B: The converse situation further down the sperm tail where in the remaining axoneme (9 + 2*) three of the doublets appear to be in the process of converting (white arrows). C: In this section the microtubule is present in a triplet-like state (inset). D: Towards the tip of the sperm the remaining axoneme only has one remaining doublet. The other 8 doublets have converted. E: Serial sectioning reveals partial interconversions of individual doublets (white arrows). F: Conversion of a doublet, via in intermediate (unfolded?) state.

## Discussion

Spermatogenesis in the acoel *Symsagittifera roscoffensis* appears to occur in a manner broadly similar to that seen in related species (Fig 1 and Fig 2) [10-12, 16, 18, 23, 32-33, 35]. We describe many aspects of this process in detail, focusing on 3 main topics: (a) the relationship between the nucleus and adjacent membranes, both the plasma membrane and the sheaths of internalised flagella, (b) the origin and fate of dense granules containing filamentous material, and (c) the conversion between microtubular forms along the tail of the spermatozoon.

### Contacts between the nucleus and other organelles

In the case of *Symsagittifera roscoffensis*, it seems likely that elongation of the nucleus, initially driven by the manchette microtubules and later by interactions with the incorporated flagella, pushes it past the mature dense granules, which fuse with it to make a thick peri-nuclear layer. Where this contacts the sperm’s plasma membrane, the nucleus forms attachments. The degree of fusion of the membrane contact site-anchored dense granules and the nuclear envelope itself cannot be assessed by TEM alone. It is possible that there is no fusion at all and we are observing two contacts, one between the plasma membrane and the Golgi-derived dense granules and a second between the dense granules and the nuclear envelope. The conformational change of the internal filaments may collapse this structure, so it appears to be a direct contact between the nuclear envelope and the plasma membrane.

We have shown that the flagella migrate into the cystosol and come to lie parallel to the nucleus. We have identified structures that might provide a mechanism for this. The free flagella represent an axoneme surrounded by a thin layer of cytoplasm and a flagella-associated plasma membrane. If formation of the flagellar sheath involved them being drawn into the cytoplasm rather than merely fusing laterally with the spermatocyte, one might imagine a mechanism by which the plasma membrane becomes associated with the flagellum at some form of collar as it is ’pulled’ in, otherwise the surface of the spermatocyte would flow over the flagellar axoneme and there would be no cytoplasmic canal formed. We suggest that the multilamellar electron-dense complex (MEC) we observe surrounding the plasma membrane of the internalising flagellum is this structure (Fig 14). It is well placed both to initiate and maintain the canal/sheath as it is drawn in. This structure is also associated with a novel membrane contact site at the junction between the base of this cytoplasmic canal and the nuclear envelope, physically linking the flagella to the nucleus. This would allow the processes of nuclear elongation and that of flagellar incorporation to proceed in unison.

A combination of both lateral fusion of flagella and subsequent formation of cytoplasmic canals (sheaths) has been suggested to give rise to the complex set of membranes observed during spermiogenesis around the internalised flagella in the closely related acoel *Paratomella rubra* [17]. Although there are significant differences between *S. roscoffensis* and *P. rubra* in terms of the final structure of the spermatozoon; these authors also identified the intimate association of the internalised flagella with the nucleus during spermiogenesis (this persists in the final spermatozoon in the latter case). The authors identify a gap in the nuclear envelope between the nucleus and the internalised flagella, and, as we see in *S. roscoffensis*, they identify microtubules in close proximity to these junctures. They do not identify a MEC or a contact site between the flagella and the nucleus, but it is possible that this is short-lived and could have been overlooked. It also underlines how certain elements of the process of spermiogenesis are common to acoels, though others may be lost or modified.

### Dense granules and formation of a thick peri-nuclear layer

Structurally complex, Golgi-derived vesicles have been observed in other, related acoels [17, 26, 30]. Dense granules arise in the Golgi and accumulate on the medial keel axoneme of the spermatids and on the manchette microtubules of the elongating nucleus. Later in development, the medial keel axoneme begins to restructure and the circle of 10 breaks into two lines or semi-circles of 5. The DGs relocate to a narrow domain of the plasma membrane in the immediate vicinity of the lateral axonemes. At this site they form a series of intimate membrane contact sites in which the DG membrane is directly opposed to the plasma membrane. The DGs do not form direct physical interactions with the lateral microtubules themselves, suggesting that the *membrane* in the vicinity of the lateral axonemes is in some way different from that covering the rest of the spermatozoon. At this site they undergo a dramatic structural reorganisation; their internal electron dense filaments (initially held in a geometric pattern by filamentous cross fibres) align perpendicular to the plasma membrane before condensing into an electron-dense core with spider’s-web like projections reaching out to the vesicular membrane.

The structural changes we see are reminiscent of the conformational changes observed in filaments during the acrosomal reaction (acrosome exocytosis) observed in other phyla during sperm maturation and fertilisation. Acrosomal fusion usually occurs between the closely apposed outer acrosomal membrane (OAM) and the plasma membrane. This results in the controlled release of the soluble protease components of the acrosome whilst an insoluble pseudo-crystalline matrix, recently shown to be an amyloid [41–45] is retained. Such amyloids (self-assembled complexes of beta-sheets) are resistant to the proteolysis characteristic of the activated acrosome environment, and potentially allow a regulated release of contents [43–46]. Other authors have noted the presence of granules in acoel spermatozoa and some have suggested that they might be components of an acrosome [10,12,14], although it is generally accepted that, with the exception of putative vesicles at the very tip of the aquasperm of some Nemertodermatida [22], acrosomes are missing from acoel sperm. Our results reveal the possibility that in some acoels the acrosome has become modified into an extended nucleus plasma-membrane contact site. It would be interesting to ascertain if any acrosome-like biochemical properties were retained.

### Microtubular structures in flux

Unlike other *Symsagittifera* spp. which have been reported to possess 8 or 12 medial keel atypical microtubules, *S. roscoffensis* has 10, which align in two groups of five that resemble musical staves (Fig 7). These microtubules have been stained in some convolutids (though not very strongly) by antibodies against beta tubulin [47, 25]; but not by antibodies raised to either alpha-tubulin or acetylated alpha-tubulin. In agreement with observations of Raikova and Justine on the microtubular system of *Convoluta saliens* [34], our results suggest that these atypical microtubules are contiguous with conventional doublets of microtubules. We have also identified what appear to be intermediate forms linking conventional doublet microtubules and the dense core microtubules. Some of these are triplet structures reminiscent of the triplet microtubules of canonical basal bodies (though arranged radially rather than linearly). The individual dense core microtubules of the staves are indistinguishable from those that constitute the centrioles of the lateral axonemes and bundle with them. We conclude that the axial keel axoneme/stave most likely represents an extended, dynamic, residual or altered basal-body, a so-called ‘centriolar derivative’.

## Conclusions

We have examined spermiogenesis in the acoel *Symsagittifera roscoffensis*; in particular, the fate of thousands of dense granules, and of the flagellar microtubules and atypical dense-cored microtubules which form the axial keel characteristic of convolute, ’aberrant’ sperm. We identify the multilaminate electron-dense complex (MEC) as an extended membrane contact site that maintains a close linkage between the nuclear envelope and the plasma membrane-derived flagellar ‘sheath’ as the developing spermatozoan undergoes structural re-organisation. The axial keel axoneme appears to be a long, elaborate centriolar-derivative, which we suggest facilitates flagellar incorporation by drawing the flagella in as it elongates. We also describe how Golgi-derived granules, by means of membrane contacts between themselves, a sub-domain of the plasma membrane and the nuclear envelope, act to ‘rivet’ the nucleus to the plasma membrane. We believe this to be the first description of an extended nucleus-plasma membrane contact site.

## Acknowledgements

This work was supported by the European Research Council (ERC-2012-AdG 322790)

